# Trees as a Reliable Carbon Capture in Urban Spaces: A Case Study in Kalaburagi

**DOI:** 10.1101/2024.07.22.604147

**Authors:** S K Shreyas, DJ Dwarka

## Abstract

The urgency of climate action has never been more apparent, and this research seeks to align the vital task of carbon mitigation with pragmatic solutions grounded in urban forestry. The objectives of this study encompass the quantification of carbon and CO_2_ stocks within the urban trees of Kalaburagi city, India, the analysis of the relative abundance of tree species, and the dissemination of findings aimed at raising awareness about the imperative of climate action. This study follows a systematic sampling approach to measure and collect data. Physical measurements were taken of each tree species spanning 20 hectares, and the readings were enumerated using allometric formulas to obtain the carbon and CO_2_ stocks (*in metric tons*). A total of over 500 individual trees belonging to 20 families were recorded. The total Biomass stood at 188.286 t, Carbon stocks reached 89.436 t, and CO_2_ stocks reached 327.871 t in the study area. *Azadirachta indica* had the highest relative abundance and sequestration potential, followed by other members of the family Meliaceae and Fabaceae, *Murraya koenigii* had the lowest carbon storage potential. The results satisfied the Shannon-Simpson indices. This research is not merely an academic endeavour; it is a call to arms, a clarion call for cities to recognize the invaluable contribution of their arboreal denizens in the struggle against climate change.

## Introduction

In an era characterized by the escalating threat of climate change, the imperative for carbon mitigation strategies has never been more compelling. The current concentration of CO_2_ in the atmosphere is 421 parts per million (*ppm*) (*NOAA*, 2023), which is double the amount of carbon during pre-industrial times at 230 ppm. This increase is attributed to human activities such as the burning of fossil fuels for energy, animal agriculture, transportation, and urbanization. Activities such as these release large amounts of greenhouse gases (GHGs), such as carbon dioxide (CO_2_), into the atmosphere. CO_2_ traps the sun’s heat and warms the planet, which has been beneficial and is the main reason for liquid water on the surface of the planet, and an increase in CO_2_ concentration has given rise to global warming.

A target to limit warming by no more than 1.5°C was given in the Paris Agreement, but countries around the world failed to meet this agreement (*Paris Agreement*, 2015). We are expected to hit 1.5°C at least once by the year 2030 and even worse, hit 2.0°C by 2050 (Solene *et al*., 2022). All climate data indicate that we are headed to the future of global boiling (*IPCC* 2023). Urban environments, often regarded as crucibles of human activity, are now emerging as battlegrounds against rising carbon dioxide (CO_2_) levels. In this context, the role of urban trees as formidable allies in carbon capture and storage has attracted unprecedented attention. (*IPCC* 2022)

Urbanization presents both opportunities and challenges. It can stimulate economic, social, and technological growth, benefiting society with improved living conditions, healthcare, and job opportunities. However, it also leads to issues like overcrowding and environmental degradation. To achieve sustainable and eco-friendly urbanization, comprehensive land use planning is essential. Urban green spaces (UGS) such as parks, gardens, and roadside vegetation must be integrated into urban settlements. UGS plays a crucial role in reducing air pollution, addressing climate change, and offering ecosystem services. In many developing countries, including India, rapid urbanization is causing significant deterioration of UGS. This review focuses on the challenges associated with creating and maintaining UGS in the Indian context, drawing from available reports highlighting problems related to poor land use planning. Challenges in UGS management and maintenance are described, with irregular watering being a major issue. The depletion of groundwater resources due to increasing water demand has led to poor watering practices, further deteriorating UGS in some Indian cities. (Manish and Avtar, 2019).

While developed countries like the USA and the UK have historic emissions, developing nations also appear to exhibit significantly high emissions. China, followed by India, has the highest emissions, primarily driven by their pursuit of becoming developed nations. Additionally, due to disparities in their stages of development, these two countries have put forth distinct types of emissions reduction commitments. China, with its relatively advanced level of industrialization and urbanization, has formally pledged to reach its peak CO_2_ emissions by 2030, to achieve this milestone as soon as possible (NDRC 2015). In contrast, India, struggling with severe electricity shortages and having very low CO_2_ emissions *per capita*, has not established a specific target for controlling its absolute emissions. Instead, it has committed to reducing emissions per unit of gross domestic product (GDP) by 33–35% by 2030 (MoEFCC). Given the substantial emissions they generate and the representative nature of their emissions reduction commitments, both China and India have significant implications for gaining a deeper understanding of CO_2_ emission growth and peak levels in the developing world.

Countries like China are employing air-capturing units that help capture carbon (DAC 2011) this promotes the dystopian ideology and rejects Urban Green spaces. Trees are natural carbon sinks; they do this by capturing CO_2_ molecules during the process of photosynthesis. Hence, they are the safest and most efficient allies in our fight against climate change. Trees also help absorb pollutants that occur in urban environments such as Polyaromatic Hydrocarbons which have adverse effects on our health and can cause respiratory ailments (Rahman *et al*., 2003).

This research endeavour embarks on a systematic exploration of the carbon capture potential of trees inhabiting urban spaces, with a particular focus on Kalaburagi City, Karnataka. The investigation delves into the biomass, carbon stocks, and relative abundance of tree species within this urban landscape. By quantifying the carbon sequestration attributes of these trees, the study aspires to shed light on their capacity to act as reliable carbon sinks (Nowak *et al*., 2013).

Some cities like Singapore which boasts itself as an urban forest have taken a step further than just planting trees, city consists of giant structures called the super trees that are an eco-architectural marvel that combine the aesthetics and skill of both architecture and Mother Nature (Meredith Davey 2011). Such hybrid approaches maybe made in metropolitan cities to add aesthetics and value to the urban areas without compromising carbon sequestration in the process. These approaches provide motivation in areas of high economic interest. A change in the policies to alleviate socioeconomic problems and create green jobs in the process, a promise that was delivered to Singapore by its governing body (*Natgeo*, 2017)

## Materials and Methods

### Study Area

The city of Kalaburagi, situated at coordinates 17.329°N and 76.825°E, was formerly known as Gulbarga. It holds significant prominence as a major city within the Hyderabad-Karnataka region, located in the northern part of the Karnataka state in India. Kalaburagi serves as the capital of the Kalaburagi District, which is one of the 31 districts in the state of Karnataka. The city is administered by the Kalaburagi Mahanagara Palike, also known as the Kalaburagi City Corporation (KCC). Covering an area of 64.00 km^2^, Kalaburagi and the entire district are situated on the Deccan Plateau, with elevations ranging from 300 to 750 meters above sea level. The prevalent soil type in the region is black soil. Kalaburagi experiences a semi-desert climate, characterized by dry conditions, with temperatures varying from 8°C to 45°C. The annual rainfall in the area averages 750mm. Although the region has historically lagged in terms of industrial development, the rapid urbanization occurring in the region is now manifesting signs of substantial industrial growth (*Wikipedia*, 2023).

**Plate 1.**
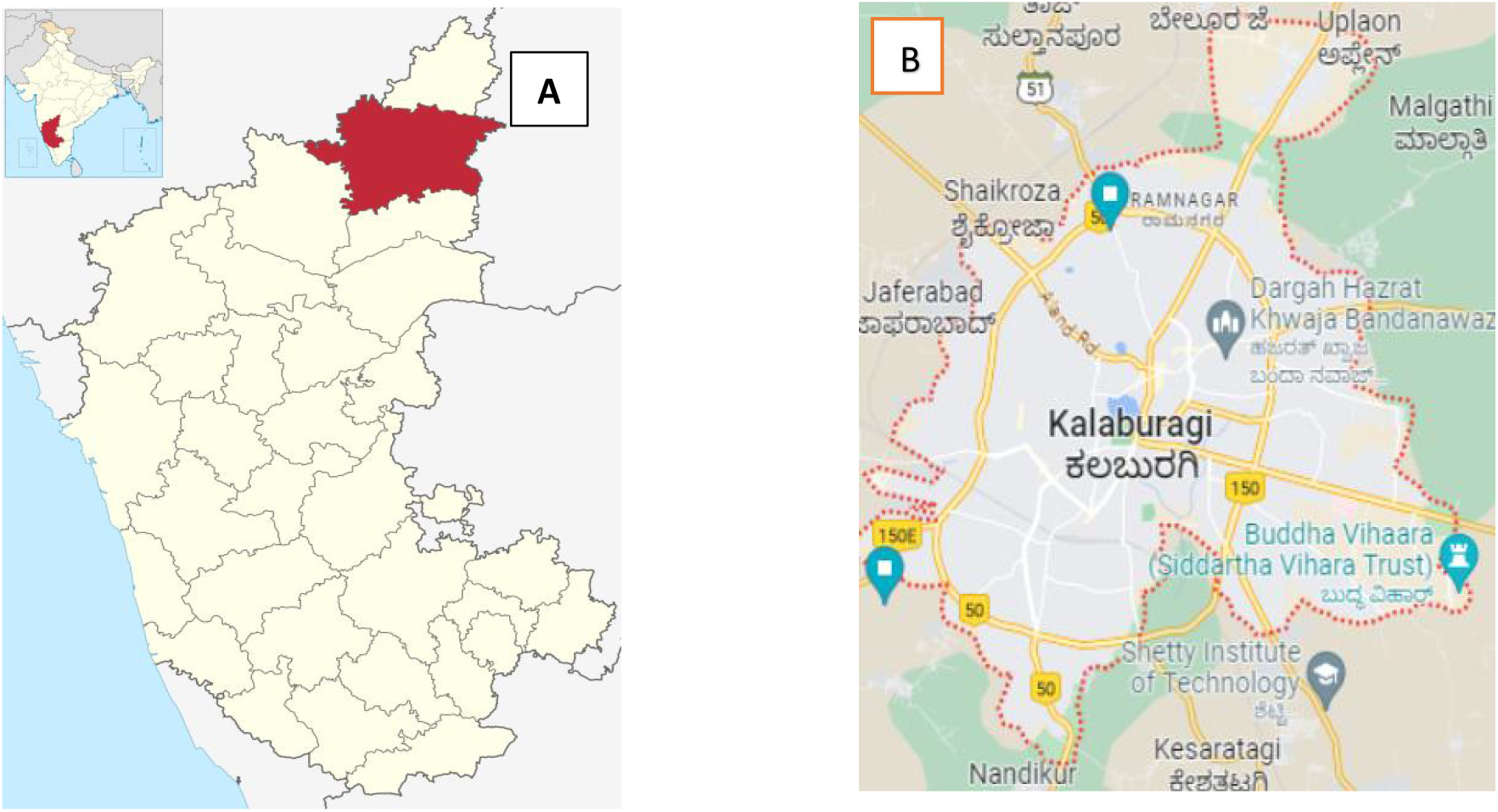
(a) Location of Kalaburagi district (in red) in the state of Karnataka, India. (b) Kalaburagi city is the governing capital of the district and spans over 64.00 km^2^. Source: Google maps.

### Methodology

This study focuses on the area inward to the city’s outer ring road. An area spanning 34.92 km^2^ was selected. Twenty plots of 100 m X 100 m (1 ha) each were selected throughout the study area, hence a total of 20 ha. A systematic tree sampling approach was preferred to maintain proper scientific conduct. The plots were taken in a randomized manner to maximize cover most of the study area.

In each plot, the number of trees was noted along with measurements of the height and girth of the tree trunk. The Diameter at Breast Height (DBH) was recorded at 1.3m from the base of the tree, and the height was measured using a clinometer and measuring tape. Data from the field survey were used to calculate the volume of the trees. The Allometric equations are mentioned below.

Above-ground biomass (ABG) was estimated by the equation:

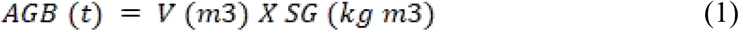

Where, V= Volume, SG = Specific gravity

The Specific gravity or wood density values (SG) of most trees were taken from Reyes *et al*. (1996) and the ICRAF Wood Density database. After this, the below-ground biomass (BGB) was estimated by multiplying the AGB by 0.26.

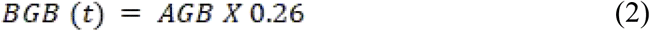

The sum of AGB and BGB gives us the Total Biomass which is then used to estimate the Carbon stocks (C). The carbon fraction value (0.475) is a default.

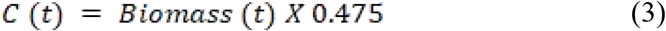

To estimate the CO_2_ Stock, the carbon stock values are multiplied by **3.666**.

The assessment of credibility involved an examination using the Shannon-Weiner and Simpson indices. Both indices are valuable tools for assessing and comparing the biodiversity in ecological communities.

The Shannon Index (H) formula is as follows:

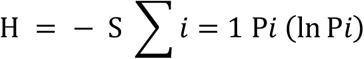

The Simpson index formula is as follows:

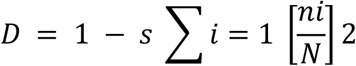

Relative Abundance of the tree species was calculated by the formula:

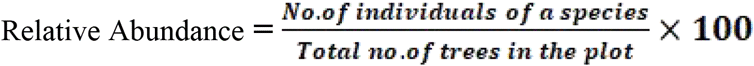

## Results and discussion

### Relative abundance

A total of 541 individual trees belonging to 20 families were recorded. *Azadirachta indica* was the dominant tree species followed by *Cocos nucifera, Peltophorum pterocarpum, Monoon longifolium* and *Milletia pinnata*. The high relative abundance of *Azadirachta indica* (24.58%) is likely due to its cultural importance in the region. It is also native to our country and hence, is better adapted to grow in the region. Among other abundant species are members of Fabaceae such as *Peltophorum pterocarpum, Milletia pinnata* and *Samanea saman*. (Table 1.). They exceptionally seem to perform well due to their ability to grow in soils with low fertility and can tolerate low rainfall. They also seem to be resilient to cattle and vandalism pressures (Rahman *et al*. 2014). Whereas the native species such as *Ficus religiosa* seem to struggle as they require protection and grow slowly when compared to non-native trees. Hence, they are not preferred by urban planners for roadside tree plantations (Ragula & Chandra. 2020).

**Table 1.** Relative Abundance of tree species in sample plots throughout the Kalaburagi study area.

| Sl. No | Tree name | Family | Number of trees in the plots | Relative Abundance (in %) |
| --- | --- | --- | --- | --- |
| 1 | *Azadirachta indica* A. Juss. | Meliaceae | 133 | 24.58 |
| 2 | *Cocos nucifera* L. | Arecaceae | 63 | 11.65 |
| 3 | *Peltophorum pterocarpum* (DC.) K. Heyne | Fabaceae | 55 | 10.17 |
| 4 | *Monoon longifolium* Sonn. B. Xue & R. M. K. Saunders | Annonaceae | 48 | 8.87 |
| 5 | *Millettia pinnata* (L.) Panigrahi | Fabaceae | 38 | 7.02 |
| 6 | *Tectona grandis* L. f. | Lamiaceae | 30 | 5.55 |
| 7 | *Samanea saman* (Jacq.) Merr | Fabaceae | 26 | 4.81 |
| 8 | *Terminalia catappa* L. | Combretaceae | 24 | 4.44 |
| 9 | *Mangifera indica* L. | Anacardiaceae | 24 | 4.44 |
| 10 | *Syzygium cumini* (L.) Skeels. | Myrtaceae | 13 | 2.40 |
| 11 | *Alstonia scholaris* (L.) R.Br. | Apocynaceae | 8 | 1.48 |
| 12 | *Dalbergia sissoo* Roxb. | Fabaceae | 6 | 1.11 |
| 13 | *Ficus religiosa* L. | Moraceae | 5 | 0.92 |
| 14 | *Cassia* sp. | Fabaceae | 5 | 0.92 |
| 15 | *Delonix regia* (Boj. ex Hook.) Raf. | Fabaceae | 5 | 0.92 |
| 16 | *Holoptelea integrifolia* (Roxb.) Planch. | Ulmaceae | 5 | 0.92 |
| 17 | *Leucaena leucocephala* (Lam.) de Wit | Fabaceae | 5 | 0.92 |
| 18 | *Moringa oleifera* L. | Moringaceae | 4 | 0.74 |
| 19 | *Vachellia nilotica* (L.) P.J.H.Hurter & Mabb. | Fabaceae | 4 | 0.74 |
| 20 | *Murraya koenigii* (L.) Sprengel | Rutaceae | 4 | 0.74 |
| 21 | *Spathodea campanulata* P.Beauv. | Bignoniaceae | 4 | 0.74 |
| 22 | *Manilkara zapota* (L.) P.Royen | Sapotaceae | 3 | 0.55 |
| 23 | *Psidium guajava* L. | Myrtaceae | 3 | 0.55 |
| 24 | *Tamarindus indicus* L. | Fabaceae | 3 | 0.55 |
| 25 | *Phyllanthus emblica* L. | Phyllanthaceae | 2 | 0.37 |
| 26 | *Ficus racemosa* L. | Moraceae | 2 | 0.37 |
| 27 | *Millingtonia hortensis* L. f. | Bignoniaceae | 2 | 0.37 |
| 28 | *Gliricidia sepium* (Jacq.) Steud. | Fabaceae | 2 | 0.37 |
| 29 | *Carica papaya* L. | Caricaceae | 2 | 0.37 |
| 30 | *Conocarpus erectus* L. | Combretaceae | 2 | 0.37 |
| 31 | *Artocarpus heterophyllus* Lam. | Moraceae | 1 | 0.18 |
| 32 | *Albizia lebbeck* (L.) Benth. | Fabaceae | 1 | 0.18 |
| 33 | *Annona squamosa* L. | Annonaceae | 1 | 0.18 |
| 34 | *Citrus × limon* (L.) Osbeck | Rutaceae | 1 | 0.18 |
| 35 | *Morinda citrifolia* L. | Rubiaceae | 1 | 0.18 |
| 36 | *Prosopis cineraria* (L.) Druce | Fabaceae | 1 | 0.18 |
| 37 | *Senna siamea* (Lam.) Irwin et Barneby | Fabaceae | 1 | 0.18 |
| 38 | *Swietenia macrophylla* King. | Meliaceae | 1 | 0.18 |
| 39 | *Thespesia populnea* (L.) Sol. ex Corrêa | Malvaceae | 1 | 0.18 |
| 40 | *Ziziphus mauritiana* Lam. | Rhamnaceae | 1 | 0.18 |
| 41 | *Ficus benjamina* L. | Moraceae | 1 | 0.18 |

**Table 2.** Location of sample plots in Kalaburagi city, and the dominant species in each plot.

| Sample Plots | Location | Total no of trees in the plot | Dominant species in the plot |
| --- | --- | --- | --- |
| P1 | Dariyapur GDA Layout | 28 | *A. indica* |
| P2 | St. John School | 18 | *A. indica* |
| P3 | PM Boys hostel | 32 | *A. indica* |
| P4 | Public Park | 57 | *P. pterocarpum* |
| P5 | Santosh colony | 25 | *M. longifolium* |
| P6 | Near PNT Quarters | 42 | *C. nucifera* |
| P7 | Rajapur | 25 | *P. pinnata* |
| P8 | Agricultural area 1.0 | 9 | *T. grandis* |
| P9 | Ashok nagar | 41 | *A. indica* |
| P10 | KHB Aland Colony | 25 | *M. longifolium* |
| P11 | Basveshwar Colony Road | 35 | *M. longifolium. A. indica* |
| P12 | Shahbazar Garden | 14 | *S. saman, A. indica* |
| P13 | Tudkur | 21 | *T. grandis* |
| P14 | Bhavani Nagar | 33 | *A. indica* |
| P15 | Godutai Nagar | 20 | *C. nucifera* |
| P16 | Manikeshwari Colony | 31 | *A. indica* |
| P17 | Islamabad Colony | 26 | *M. indica* |
| P18 | Khaja Colony | 21 | *A. indica* |
| P19 | Agricultural area 2.0 | 8 | *A. indica* |
| P20 | Central Main road | 30 | *P. pterocarpum* |

**Table 3.**
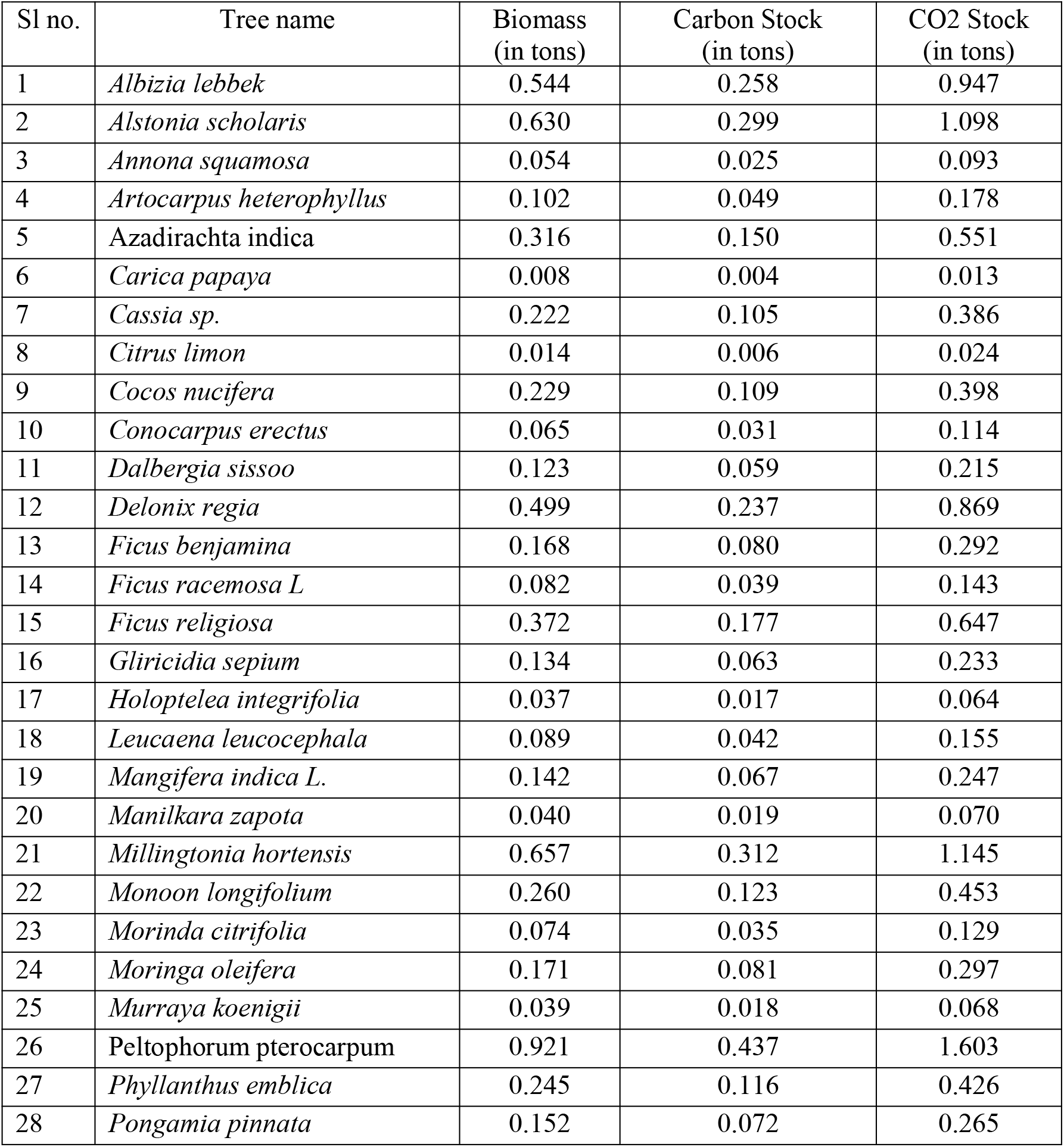
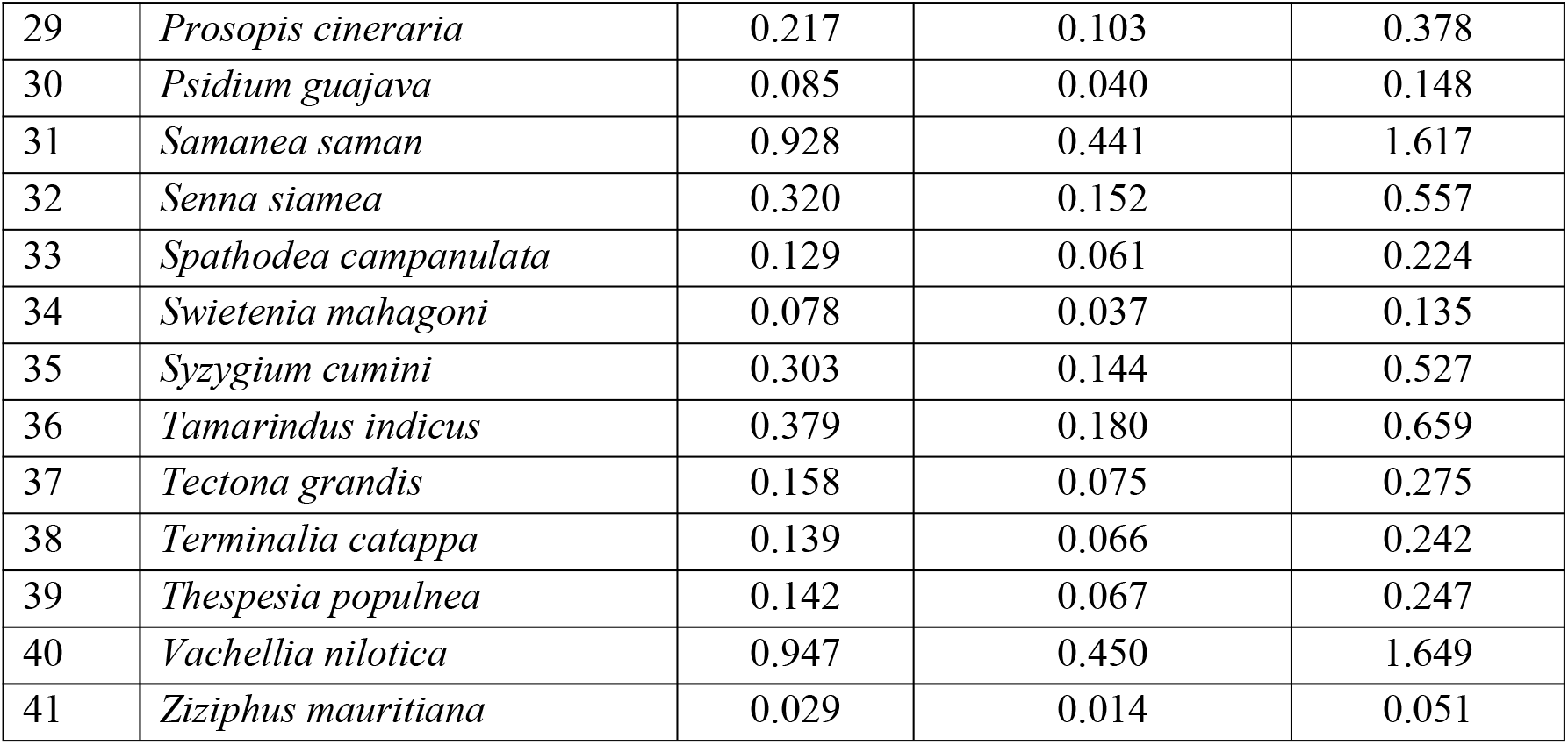
Species-wise Average biomass, carbon capture and CO_2_ Stock.

Trees from the Fabaceae family have been employed in afforestation efforts throughout Southeast Asia and are now commonly seen in urban areas across the region. Although they exhibit rapid growth, they can also display a certain level of invasiveness, often being non-native and competing with indigenous species for resources. Conversely, they may thrive in challenging environmental conditions and the presence of various disturbances (Rahman *et al*. 2014).

Plot-level data indicates an average of 27 trees per plot within a set of 20 one-hectare plots. Among these, plot P4 stood out with the highest tree count among the sampled plots. Similarly, several plots, namely P3, P4, P6, P9, P11, P14, and P16, boasted more than 30 trees per plot. *Azadirachta indica* dominated plots P1, P2, P3, P9, P11, P12, P14, P16, P18, and P19, while *Monoon longifolium* was the dominant species in P5, P10, and P11. *Peltophorum pterocarpum* claimed dominance in plots P4 and P20, while *Cocos nucifera* held sway in P6 and P15. *Tectona grandis* dominated in P9 and P13, while *Mangifera indica* took precedence in P17. In residential areas, trees flourished, benefiting from the nurturing care and protection provided by the residents. In contrast, plots P8 and P19, situated within the urban fabric of the city, were primarily earmarked for agricultural use and thus exhibited a limited tree population. This was due to a substantial portion of the land being allocated for agricultural practices such as crop cultivation. However, the city’s swift urbanization has triggered a transformation of these fields into residential and civic accommodations, subsequently altering the local landscape.

This research discovered that tree species characterized by broader canopy coverage, such as *Samanea saman, Ficus religiosa, Acacia nilotica* and *Terminalia catappa*, exhibited excellent performance in various settings, except roadside plantations. These species demonstrated a significant capacity to capture substantial amounts of carbon (Sala *et al*., 2012). In contrast, *Monoon longifolium* effectively leveraged its ability to grow upright, making it well-suited for roadside environments.

The study also found that socio-economic factors, such as poverty and living conditions, had a significant impact on the distribution of trees and their potential for carbon sequestration. Regions with affluent communities demonstrated greater care for trees, evident through the higher abundance and volume of trees in those areas.

Conversely, communities residing in slums showed less interest in tree planting and protection, likely due to their lack of engagement with the subject. While this situation may have negative implications, it underscores that increasing urbanization exacerbates existing inequalities. It is unjust for these communities to endure conditions like tightly packed housing, narrow streets, stagnant drainage systems, and poor ventilation without the benefits of trees to help filter some of the pollution. This could further worsen issues related to disease transmission and impact the overall health of these communities (Ramaiah and Avtar, 2019).

### Carbon and CO_2_ Stocks in trees

The ten tree species that exhibited the highest carbon sequestration capabilities were *Azadirachta indica, Albizia lebbek, Alstonia scholaris, Delonix regia, Ficus religiosa, Samanea saman, Peltophorum pterocarpum, Millingtonia hortensis, Tamarindus indicus*, and *Vachellia nilotica*. On the contrary, *Ziziphus mauritiana* and *Murraya koenigii* showed the lowest carbon capture abilities. Given that this study encompassed diverse urban areas, including bustling main roads and residential neighbourhoods, the results offer comprehensive insights into the carbon capture potential of urban trees in the city. The combined volume of the sampled trees amounted to 248.358 m^3^, with a gross biomass of 188.286 metric tons. The total carbon stocks reached 89.436 metric tons, and CO2 Stocks reached 327.871 metric tons. These findings align with the results of previous researchers such as Yin *et al*.(2012), Bohre *et al*.(2012), & Ragula and Chandra (2020). The area covering 20 hectares has a Simpson and Weiner index value of 2.72, and there is a similarity in the Shannon index value of 0.108 when compared to the findings of Baragpur *et al*. in 2022.

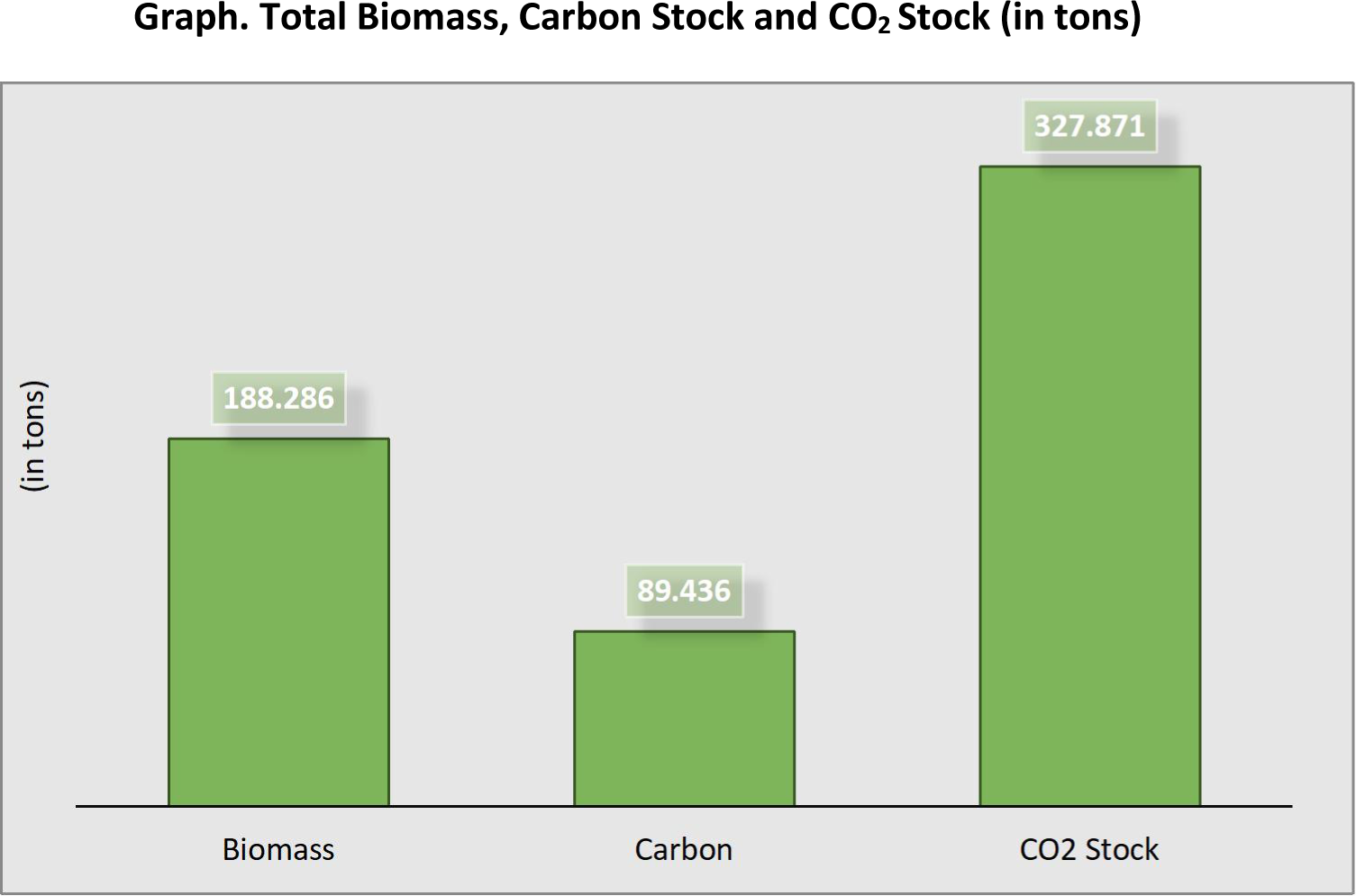

While most studies emphasize the importance of planting native species to promote and preserve local biodiversity (Ragula and Chandra, 2020), it is essential to recognize that urban spaces within a city represent highly artificial conditions. Even native species may struggle to adapt to these environments. This study advocates for a hybrid approach that incorporates a diverse selection of trees, both native and exotic, to harness the benefits each type offers. Native species can prevent the homogenization of tree populations, while exotic species, such as members of the Fabaceae family, can yield satisfactory results due to their ability to thrive rapidly in urban settings, facilitating the capture of substantial amounts of carbon from the atmosphere. The increasing need for urban green spaces is crucial to mitigate the ‘Urban Island Effect,’ where cities trap and retain more heat due to roads and buildings, exacerbating the impacts of climate change (Yang et al., 2016).

## Conclusion

In conclusion, this research has provided valuable insights into the distribution and performance of tree species in our urban environment. *Azadirachta indica*, owing to its cultural significance and native adaptability, emerged as the dominant species, while members of the Fabaceae family showcased exceptional resilience and growth in adverse conditions. The study highlighted the pivotal role of socio-economic factors in shaping tree distribution patterns, with affluent communities demonstrating greater care for trees compared to slum areas. This underscores the need for equitable urban planning to ensure all communities benefit from the positive effects of trees. Furthermore, the research emphasized the importance of a hybrid approach to tree planting, incorporating both native and exotic species. While native species play a vital role in preserving local biodiversity, exotic species, especially those from the Fabaceae family, offer rapid growth and effective carbon capture capabilities in urban settings. The carbon sequestration potential of urban trees was also assessed, with several species demonstrating significant abilities to capture and store carbon. These findings underline the importance of urban green spaces in mitigating the effects of climate change and the urban heat island effect. In summary, this research contributes to our understanding of urban tree dynamics and calls for a balanced approach to urban forestry that considers both ecological and socio-economic factors. By doing so, we can create healthier and more sustainable urban environments for all residents.

## Notes

### Competing Interest Statement

The authors have declared no competing interest.

